# Untargeted metabolomic analysis in naturally occurring canine diabetes mellitus identifies similarities to human Type 1 Diabetes

**DOI:** 10.1101/139113

**Authors:** Allison L. O’Kell, Timothy J. Garrett, Clive Wasserfall, Mark A. Atkinson

## Abstract

While predominant as a disease entity, knowledge voids exist regarding the pathogenesis of canine diabetes. To test the hypothesis that diabetic dogs have similar metabolomic perturbations to humans with type 1 diabetes (T1D), we analyzed serum metabolomic profiles of breed- and body weight-matched, diabetic (n=6) and healthy (n=6) dogs by liquid chromatography-mass spectrometry (LC-MS) profiling. We report distinct clustering of diabetic and control groups based on heat map analysis of known and unknown metabolites. Random forest classification identified 5/6 dogs per group correctly with overall out of bag error rate=16.7%. Diabetic dogs demonstrated significant upregulation of glycolysis/gluconeogenesis intermediates (e.g., glucose/fructose, C_6_H_12_O_6_, keto-hexose, deoxy-hexose, (P<0.01)), with significant downregulation of tryptophan metabolism metabolites (e.g., picolinic acid, indoxyl sulfate, anthranilate, (P<0.01)). Multiple amino acids (AA), AA metabolites, and bile acids were also significantly lower in diabetic versus healthy dogs (P<0.05) with the exception of the branched chain AA valine, which was elevated in diabetic animals (P<0.05). Metabolomic profiles in diabetic versus healthy dogs shared similarities with those reported in human T1D (e.g., alterations in glycolysis/gluconeogensis metabolites, bile acids, and elevated branched chain AA). Further studies are warranted to evaluate the utility of canine diabetes to provide novel mechanistic insights to the human disorder.

**Abbreviations:**

AA: (amino acid)
AAb: (autoantibody)
FA: (fatty acid)
HILIC: (hydrophilic interaction liquid interaction chromatography)
LC-MS: (liquid chromatography mass spectrometry)
OOB: (out of bag)
T1D: (type 1 diabetes)
T2D: (type 2 diabetes)
UF: (University of Florida)

---

Type 1 diabetes (T1D) is characterized by insulin deficiency and resulting dysglycemia^1^. Recently, metabolomic analyses have been used to study metabolic changes in T1D patients prior to autoantibody (AAb) development^2^, associated with AAb seroconversion^3^, as well as with symptomatic T1D and glycemic control^4–6^. Specifically, metabolomic alterations documented prior to AAb development in children that later developed T1D include decreased succinic acid, phosphatidylcholine, and citric acid at birth alongside increased pro-inflammatory lysophosphatidylcholine months prior to seroconversion^2^. AAb positivity is associated with low methionine and hydroxyproline, with high odd-chain triglycerides and polyunsaturated fatty acids (FA) containing phospholipids^3^. Beyond this, T1D patients with poor glycemic control demonstrate decreased glycolytic metabolites and elevated carbohydrate metabolites, branched chain amino acids (AA), short chain FA, and ketoacids^4^. Interestingly, many of these metabolic perturbations are also present in T1D patients with good glycemic control^4^.

Canine diabetes has multiple parallels to human T1D, including a requirement for lifelong insulin therapy in most cases, the development of ketoacidosis at diagnosis or during therapy^7^, and a suspected role for autoimmunity^7, 8^. Increased levels of glucose, triglycerides, non-esterified FA, cholesterol, ketones, acetate, and β-hydroxybutyrate have been reported in diabetic dogs^9^. Further research is required to determine the potential for canines with diabetes to be used as an alternative animal model of human T1D^10^, and if metabolomic analysis may identify novel biomarkers of the disease. While metabolomic profiles have been studied in healthy dogs^11, 12^ and in several disease states, including inflammatory bowel disease^13^ and degenerative valvular disease^14^, information from an untargeted metabolomics assessment of diabetic dogs is lacking. With this study, we evaluated the metabolomic profiles of fasted diabetic versus healthy control dogs, and hypothesized that diabetic dogs have metabolomic perturbations similar to those reported in human T1D patients, including alterations in carbohydrates, branched-chain AA and FA.

## Materials and Methods

### Study approval and enrollment

The study was approved by the University of Florida (UF) Institutional Animal Care and Use Committee (#201609360) and the Veterinary Hospital Research Review Committee. All dog owners provided written informed consent prior to study enrollment. Dogs with naturally occurring diabetes (n=6) and control dogs (n=6) were recruited from the hospital population at the UF Small Animal Hospital. Diabetes was diagnosed prior to study enrollment based on clinical signs (polyuria, polydipsia) in combination with persistent hyperglycemia and glucosuria. Diabetic dogs were included if they were >1 year of age with body weight >5 kg, were fasted for a minimum of 12 hours, and if female, were spayed prior to diagnosis of diabetes. Control dogs were deemed healthy based on history and physical exam findings. Control dogs were breed matched to diabetic dogs when possible, and were included if they were >1 year of age with body weight >5 kg, received no medications other than monthly flea/tick/heartworm preventatives, and were fasted for a minimum of 12 hours. All dogs were housed with their owners aside from their visit to the Small Animal Hospital.

### Sample collection

Time of blood collection was based on patient availability, but all blood samples were collected between 8am and 4 pm. Blood was collected using a needle and syringe by routine venipuncture and allotted into red top vacutainer tubes containing clot activator. Serum was separated within 30 min of collection and frozen immediately at −80°C.

### Metabolite extraction

Metabolite extraction and analysis from serum was performed as previously described^15, 16^. Briefly, 100 µL serum aliquots were prepared, and 800 µL of 8:1:1 acetonitrile:methanol:acetone was added to precipitate the proteins. The samples were allowed to cool on ice for 30 minutes before centrifugation to pellet the protein. The supernatant (750 µL) was transferred to a new microcentrifuge tube and dried under a gentle stream of nitrogen. The dried sample was reconstituted in 100 µL of 0.1% formic acid in water for liquid chromatography-mass spectrometry (LC-MS) analysis. LC-MS profiling was performed using a Thermo Q-Exactive Orbitrap mass spectrometer (mass resolution 35,000 at m/z 200) with Dionex UHPLC and autosampler. Samples were analyzed in a positive and negative heated electrospray ionization as separate injections.

### Metabolite analysis

MZmine^17^ was used to align, gap-fill, and identify metabolites across the sample set. Known metabolites were identified by matching to an internal retention time library, except in the case of lipids which were identified by fragmentation pathways from MS/MS. Random forest was used to evaluate the classification performance of metabolomics. Analyses were performed using Metaboanalyst 3.0, an open source R-based program specifically designed for metabolomics^18^. Values not present in 80% of the data were removed from analysis. Missing values were imputed using k-nearest neighbor, and the data was interquartile range filtered, sum normalized, log2 transformed and autoscaled. All statistical analyses were performed on the combined positive ion and negative ion data sets.

### Statistics

Patient characteristics (Table 1) were compared using Mann-Whitney or Fisher's exact test via GraphPad Prism software v6. Metabolites were compared across the two groups via t-test, and a heatmap of significantly different features was generated to identify clustering metabolites P<0.05 was considered significant.

**Table 1.** Patient Characteristics

| Group | Breed | Age (years) | Median Age (years) | Weight (kg) | Median Weight (kg) | Sex<sup>a</sup> | Duration since diagnosis of diabetes (months) | Median Disease Duration (months) |
| --- | --- | --- | --- | --- | --- | --- | --- | --- |
| Diabetic | Dachshund | 6.5 | 9.75 | 7.9 | 18.8 | SF | 1.5 | 4.75 |
|  | Dachshund | 13 |  | 6.9 |  | SF | 1 |  |
|  | Miniature schnauzer | 9 |  | 9.5 |  | NM | 8 |  |
|  | Labrador retriever | 13.5 |  | 28.4 |  | SF | 11 |  |
|  | Labrador retriever | 10.5 |  | 32.5 |  | NM | 11 |  |
|  | Mix (Labrador retriever mix) | 1.3 |  | 28.1 |  | NM | 1 |  |
| Healthy | Dachshund | 4 | 5.5 | 8.8 | 19.75 | SF | N/A | N/A |
|  | Dachshund | 11 |  | 5.2 |  | SF | N/A |  |
|  | Miniature schnauzer | 2 |  | 6.6 |  | SF | N/A |  |
|  | Labrador retriever | 7 |  | 31.8 |  | SF | N/A |  |
|  | Labrador retriever | 4 |  | 43.2 |  | NM | N/A |  |
|  | Flat coated retriever | 8 |  | 30.7 |  | NM | N/A |  |

<sup>a</sup>SF=spayed female; NM= neutered male

## Results

Age (P=0.29), body weight (P=0.99) and sex distribution (P=1.0) were comparable for diabetic and control groups (Table 1). For diabetic dogs, the median duration from diagnosis was 4.75 months (range: 1-11 months). Controls were breed-matched to diabetic dogs in all but one case (Table 1).

Two hundred known metabolites were detected in positive and negative ion modes from the extracted serum samples. A heat map of known metabolites that differed significantly between diabetic and healthy dogs indicated clustering of the two cohorts (Figure 1A). A similar analysis of all detected (unknown and known) metabolites not only enabled classifications based on the entire metabolome, but also improved clustering (Figure 1B). The top 50 metabolites that differed significantly between groups are shown in Figure 1B. These data demonstrate the ability of metabolomics to classify diabetic and non-diabetic canines.

**Figure 1.**
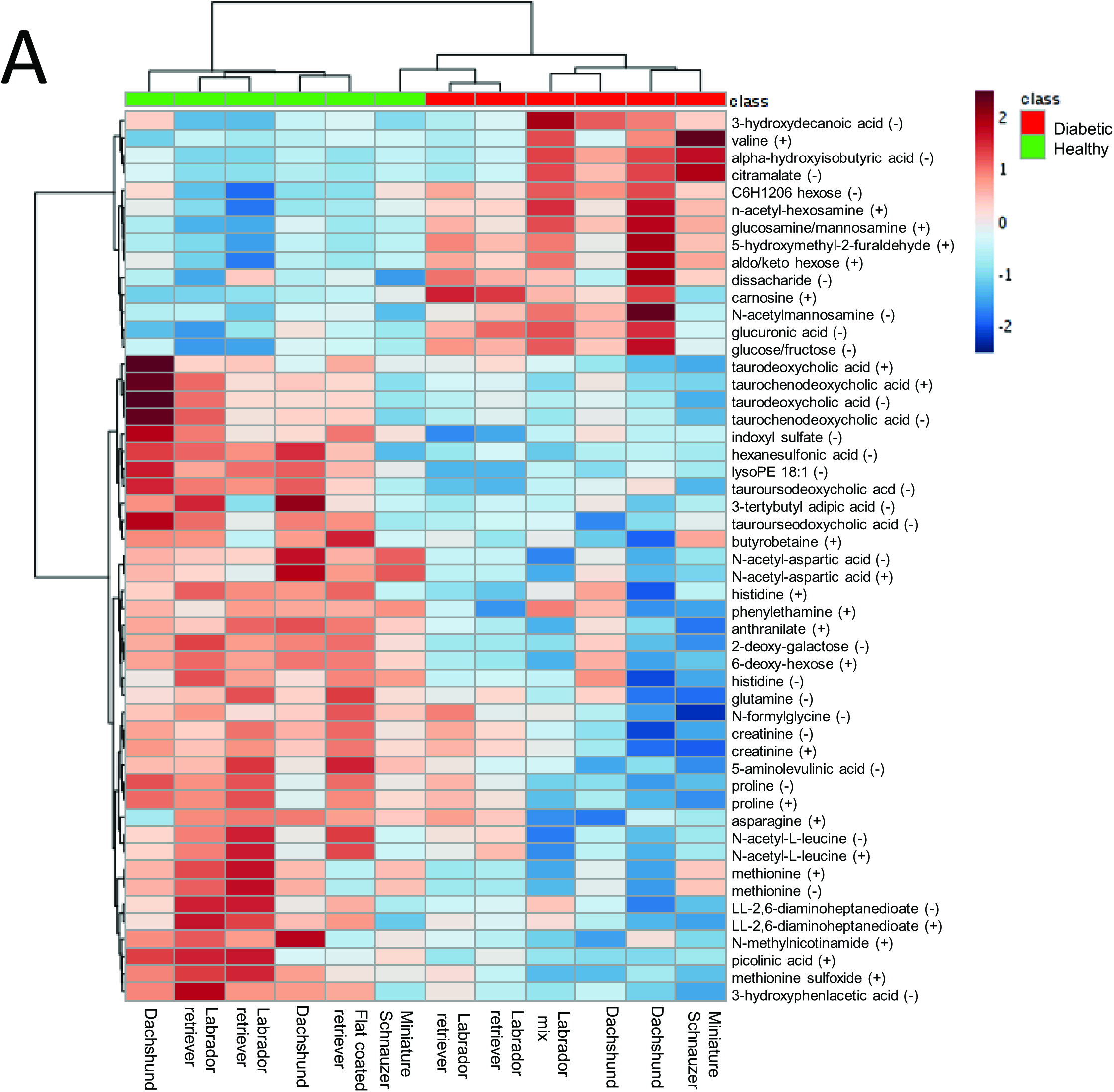
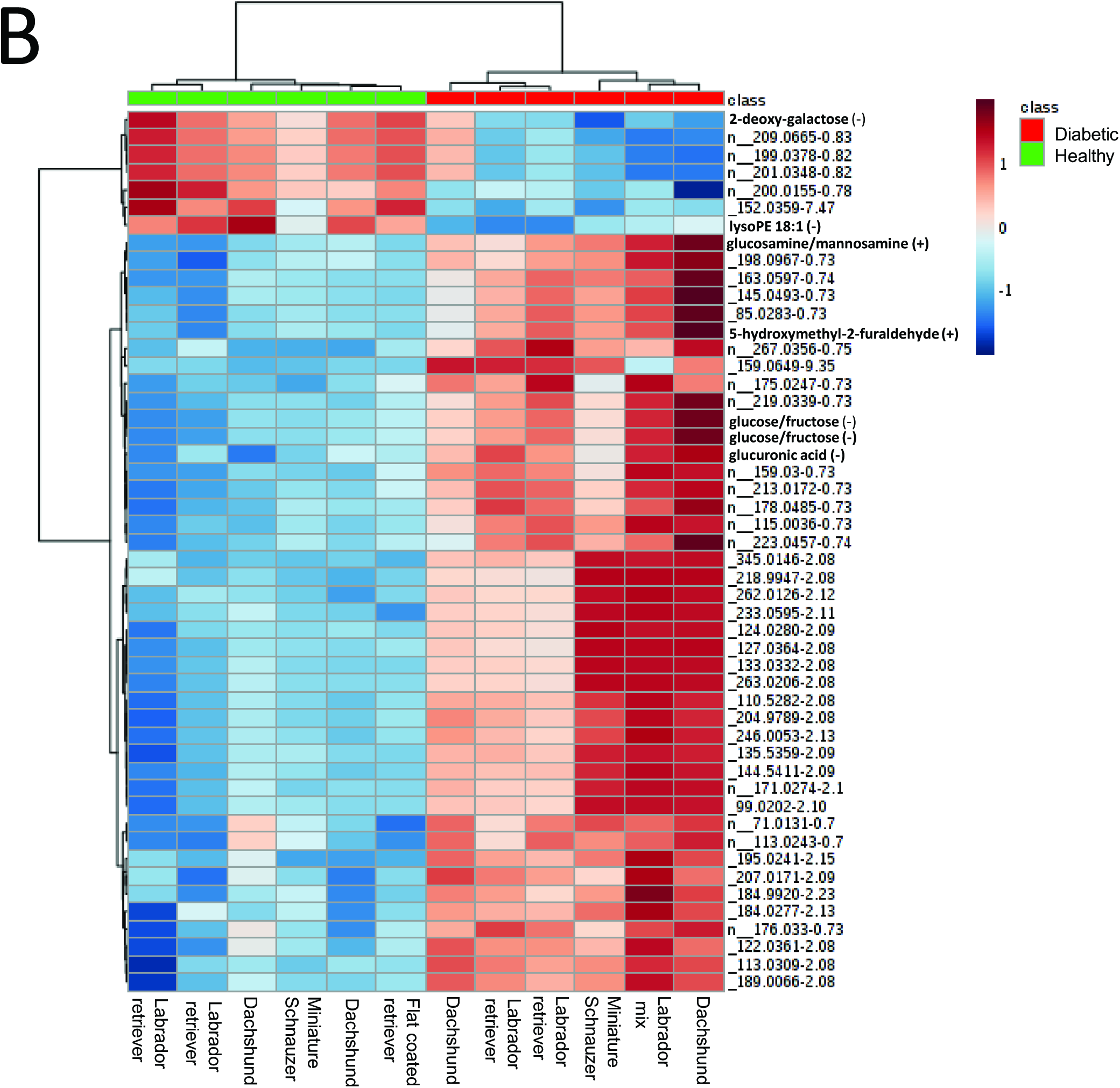
Heat map representing (A) known metabolites significantly (p<0.05) different between diabetic and healthy control dog groups and (B) top 50 known and unknown metabolites significantly (p<0.05) different between groups. Group is indicated at the top of the figure by red (diabetic, n=6) or green (healthy, n=6). Individual dog breed corresponding to column is indicated at the bottom of the figure. Data was sum normalized, log transformed and autoscaled.

The known metabolites identified are categorized by class and/or pathway and direction of change in diabetic animals as compared to controls (Table 2). As expected, glycolysis/gluconeogenesis intermediates and sugars were significantly upregulated in the diabetic group, while concentrations of metabolites involved in tryptophan metabolism were downregulated (Table 2). Multiple AA and AA metabolites were significantly lower in the diabetic group, including proline, methionine, histidine, glutamine, and asparagine, while the branched chain AA, valine, was higher. The primary and secondary bile acids, taurochenodeoxycholic acid, taurodeoxycholic acid and tauroursodeoxycholic acid, were significantly lower in diabetic dogs. The lipids lysophosphatidylethanolamine and butyrobetaine were lower in the diabetic group, while 3-hydroxydecanoic acid was higher versus control dogs.

**Table 2.** Significant Metabolites and Pathways

| **Class/Pathway and Metabolites** | **<sup>a</sup>Diabetic dogs** | **<sup>a</sup>Healthy dogs** | **p-value** |
| --- | --- | --- | --- |
| *Glycolysis/gluconeogenesis intermediates* |  |  |  |
| Glucose/Fructose | ↑ | ↓ | 0.001205 |
| Glucose/Fructose (Cl adduct) | ↑ | ↓ | 0.001314 |
| *Aminosugar and nucleotide sugar metabolism* |  | ↓ |  |
| Glucuronic acid | ↑ | ↓ | 0.00078 |
| Glucosamine/mannosamine | ↑ | ↓ | 0.000371 |
| *Sugars* |  |  |  |
| C<sub>6</sub>H<sub>12</sub>O<sub>6</sub> | ↑ | ↓ | 0.002526 |
| N-acetyl-hexosamine | ↑ | ↓ | 0.0003124 |
| Keto-hexose | ↑ | ↓ | 0.000148 |
| Deoxy-hexose | ↓ | ↑ | 0.000615 |
| 2-deoxy-D-galactose | ↓ | ↑ | 0.000369 |
| Disaccharide | ↑ | ↓ | 0.016432 |
| *Dipeptide* |  |  |  |
| Carnosine | ↑ | ↓ | 0.007063 |
| *Tryptophan metabolism* |  |  |  |
| Picolinic acid | ↓ | ↑ | 0.004321 |
| Indoxyl sulfate | ↓ | ↑ | 0.004485 |
| Anthranilate (2-aminobenzoic acid) | ↓ | ↑ | 0.000553 |
| *Organic acid and derivatives* |  |  |  |
| 5-aminolevulinic acid | ↓ | ↑ | 0.0002224 |
| Alpha-hydroxyisobutyric acid | ↑ | ↓ | 0.009782 |
| *Amino Acids and Metabolites* |  |  |  |
| Proline | ↓ | ↑ | 0.007627 |
| N-acetyl-L-aspartic acid | ↓ | ↑ | 0.000794 |
| N-acetyl-L-aspartic acid (positive ion) | ↓ | ↑ | 0.001863 |
| Methionine | ↓ | ↑ | 0.012396 |
| Methionine sulfoxide | ↓ | ↑ | 0.001815 |
| Valine | ↑ | ↓ | 0.012396 |
| Histidine | ↓ | ↑ | 0.016368 |
| Glutamine | ↓ | ↑ | 0.020218 |
| Creatinine | ↓ | ↑ | 0.021156 |
| Asparagine | ↓ | ↑ | 0.049261 |
| N-acetyl-l-leucine | ↓ | ↑ | 0.022766 |
| 3-hydroxyphenylacetic acid | ↓ | ↑ | 0.006011 |
| N-formylglycine | ↓ | ↑ | 0.040731 |
| *Bile acids* |  |  |  |
| Taurodeoxycholic acid | ↓ | ↑ | 0.035656 |
| Taurochenodeoxycholic acid | ↓ | ↑ | 0.043389 |
| Tauroursodeoxycholic acid | ↓ | ↑ | 0.005472 |
| *Organoheterocyclic compounds* |  |  |  |
| N-methylnicotinamide | ↓ | ↑ | 0.007934 |
| *Aldehyde* |  |  |  |
| 5-hydroxymethyl-2-furaldehyde | ↑ | ↓ | 0.000515 |
| *Benzenoid* |  |  |  |
| Phenethylamine | ↓ | ↑ | 0.028163 |
| *Lipids* |  |  |  |
| LysoPE 18:1 | ↓ | ↑ | 0.000226 |
| 3-hydroxydecanoic acid | ↑ | ↓ | 0.02393 |
| Butyrobetaine | ↓ | ↑ | 0.047759 |
| *Microbial origin* |  |  |  |
| Citramalate (fatty acid) | ↑ | ↓ | 0.013703 |
| LL-2,6 diaminoheptanedioate (Diaminopimelic acid)<br>(amino acid) | ↓ | ↑ | 0.049682 |
| *Other Exogenous* |  |  |  |
| Tertbutyl adipic acid | ↓ | ↑ | 0.048648 |
| Hexanesulfonic acid | ↓ | ↑ | 0.011531 |

<sup>a</sup> ↓ indicates downregulation; ↑ indicates upregulation

Random forest classification showed excellent prediction of group, with 5/6 animals from each group identified correctly and a 16.7% out of bag (OOB) error rate (Table 3). The healthy control dog misclassified to the diabetic group by this method was confirmed to be normoglycemic in both a fasted and fed state 3 months following study enrollment. The random forest importance plot identified 15 metabolites key in classifying the data, with anthranilate, glucose, glucosamine, and lysophosphatidylethanolamine having the most influence in classification (Figure 2).

**Figure 2.**
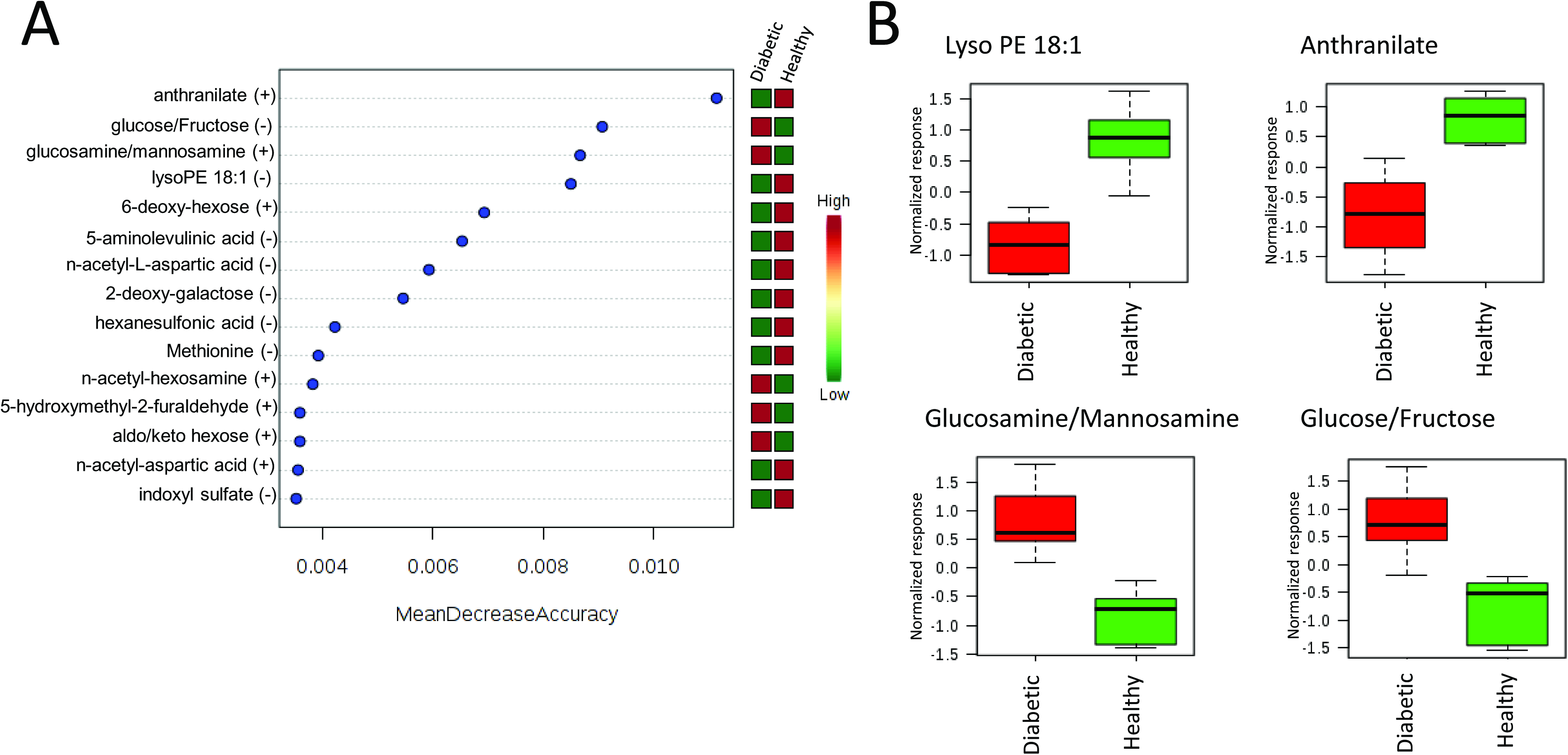
(A) Random forest variable importance plot. Mean decrease accuracy is the measure of the performance of the model without each metabolite. A higher value indicates the importance of that metabolite in predicting group (diabetic vs. healthy). Removal of that metabolite causes the model to lose accuracy in prediction. (B) Box and whisker plots of the top four metabolites from A. Data was sum normalized, log transformed and autoscaled.

**Table 3.**
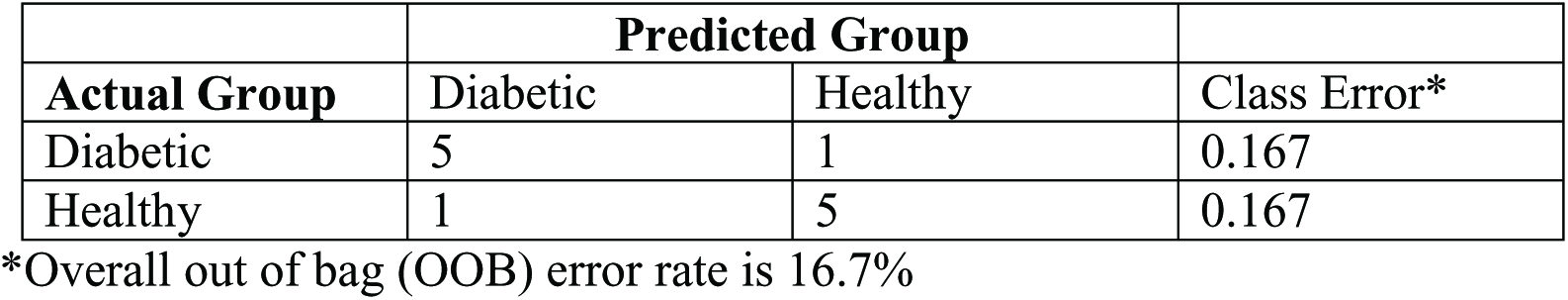
Random Forest Classification into Healthy or Diabetic Dog Groups

|  | **Predicted Group** |  |  |
| --- | --- | --- | --- |
| **Actual Group** | Diabetic | Healthy | Class Error* |
| Diabetic | 5 | 1 | 0.167 |
| Healthy | 1 | 5 | 0.167 |

\*Overall out of bag (OOB) error rate is 16.7%

## Discussion

In this study using an untargeted metabolomic evaluation, we found that the metabolomic profiles differed significantly among fasted diabetic and healthy dogs, with clear clustering of the groups using known and unknown metabolite expression. Based on random forest classification, 5/6 (83%) dogs in each group were classified correctly, an excellent result given the small sample size. The healthy dog that was misclassified as a diabetic dog (Miniature Schnauzer, Figure 1) in this analysis was confirmed to be normoglycemic 3 months following the study. This dog displayed upregulated carnosine and downregulated histidine, n-actetyl-L-leucine, and multiple bile acids (similar to diabetic dogs). Possible reasons for this discrepancy include individual variability in metabolomic profiles, environmental or dietary factors, a yet unidentified pre-diabetic state, or other undiagnosed underlying disease.

Studies examining inter- and intra-individual variability in metabolomic profiles in dogs indicate that dog breed is an important factor in metabolomic profile variance in the non-controlled (i.e. home) environment, along with some influence of gender and body weight^11, 19^. Importantly, in this study, all but two dogs were breed and gender matched and the non-breed matched dogs were body weight matched. The results of this pilot study support future research to expand the scope by increasing the number of dogs examined and implementing a longitudinal design with multiple time points. Moreover, the analytical methods employed have excellent mass accuracy (5 ppm or better) and retention time reproducibility (less than 0.1 min fluctuations), but do not separate all the sugar related species. We also plan to conduct additional studies to identify unknown metabolites with the addition of hydrophilic interaction liquid interaction chromatography (HILIC) to improve separation of sugar related metabolites.

Observed elevations in carbohydrate metabolites and alterations in glycolysis/gluconeogensis metabolites in diabetic dogs were expected and similar to humans with T1D^4^. Similarly, the branched chain AA valine was significantly elevated in diabetic dogs; branched chain AA (valine, leucine, isoleucine) have also been documented to be elevated in humans prior to and after onset of both T1D and type 2 diabetes (T2D)^4, 20^ as well as in diabetic NOD mice^21^. However, we report lower N-acetyl-L-leucine in diabetic dogs versus controls while isoleucine was not identified as significantly differing between the groups, representing a potential divergence from human T1D or simply a limitation of the small sample size. The AA methionine was lower in diabetic dogs, similar to children who develop islet AAb at an early age^3^ but in contrast to human T1D patients with poor glycemic control and reported elevations in this AA^4^. Several metabolites in involved in tryptophan metabolism (e.g., anthranilate/2-aminobenzoic acid, picolinic acid, indoxyl sulfate) were lower in diabetic dogs versus healthy controls. This contrasts to reports of increased anthranilic acid in human T1D patients compared to T2D patients and controls without diabetes^22^. Further investigation is needed to determine the implications of these findings in dogs, particularly given the importance of anthranilate in the random forest variable importance plot (Figure 2A).

Another interesting finding was the upregulation of carnosine in diabetic dogs compared with control dogs. Carnosine has been shown to inhibit the formation of some advanced glycation end products (AGEs) *in vitro*^23^ and to inhibit glucose-induced production of type IV collagen and fibronectin by podocytes in the kidneys^24^. A genetic mutation resulting reduced expression of the carnosinase enzyme that degrades carnosine protects against the development of diabetic nephropathy in human patients^24^. While dogs with experimentally induced and naturally occurring diabetes do develop renal glomerular lesions consistent with diabetic nephropathy^25, 26^, dogs with naturally occurring diabetes do not seem to commonly develop clinically significant diabetic nephropathy^27, 28^. Although this is speculated to be due to the shorter lifespan of dogs and lack of time for significant renal disease pathology to develop^27^, the elevation in carnosine could be relevant. Further study is required to confirm these findings and any relationship to the formation of AGEs in dogs.

Primary and secondary bile acid levels were found to be lower in diabetic dogs, and perturbations in a variety of bile acids have been identified in humans with T1D and T2D^4, 29^. Bile acids have been recognized as having endocrine functions and are involved in glucose homeostasis^30^. Adding to the complexity of bile acid metabolic function is the interplay with the gut microbiota^31^, which has recently been shown to be altered in humans and rodents with diabetes^32^. In the present study, gut microbiota was not evaluated and the cause of the alterations in bile acids is unknown. Given the important role bile acids play in metabolism, additional investigation is warranted.

In sum, we found that the metabolomic profiles of diabetic dogs differ significantly from healthy non-diabetic dogs. Both similarities to and differences from metabolic perturbations in humans with diabetes were identified. Further research is needed with increased sample size in a longitudinal study design and potentially, with a standardized diet to confirm the findings reported herein and identify the underlying mechanisms that contribute to these metabolic changes. Many unknowns still exist with respect to the pathogenesis of canine diabetes, and metabolomic alterations may represent important potential biomarkers to detect diabetes in dogs prior to clinical onset. Indeed, the ability to detect the pre-diabetic state may represent a key component to future discovery in this area^10^ and to further define the utility of the dog as a potential model of human T1D.

## Author Contributions

ALO conceived of the study, collected the data, analyzed and interpreted the data, and wrote the manuscript; TJG conducted the analytical measurements, performed statistical analysis, and analyzed and interpreted the data, and wrote the manuscript. CW conceived of the study, analyzed and interpreted the data, contributed to the discussion, and reviewed/edited the manuscript; MAA conceived of the study, contributed to discussion and reviewed/edited the manuscript.

## References

1 Atkinson, M. A., Eisenbarth, G. S. & Michels, A. W. Type 1 diabetes. Lancet 383, 69–82, doi:10.1016/S0140-6736(13)60591-7S0140-6736(13)60591-7 [pii] (2014).

2 Oresic, M. et al. Dysregulation of lipid and amino acid metabolism precedes islet autoimmunity in children who later progress to type 1 diabetes. J Exp Med 205, 2975–2984, doi:jem.20081800 [pii] 10.1084/jem.20081800 (2008).

3 Pflueger, M. et al. Age- and islet autoimmunity-associated differences in amino acid and lipid metabolites in children at risk for type 1 diabetes. Diabetes 60, 2740–2747, doi:60/11/2740 [pii] 10.2337/db10-1652 (2011).

4 Dutta, T. et al. Impact of Long-Term Poor and Good Glycemic Control on Metabolomics Alterations in Type 1 Diabetic People. J Clin Endocrinol Metab 101, 1023–1033, doi:10.1210/jc.2015-2640 (2016).

5 Lanza, I. R. et al. Quantitative metabolomics by H-NMR and LC-MS/MS confirms altered metabolic pathways in diabetes. PLoS One 5, e10538, doi:10.1371/journal.pone.0010538 (2010).

6 Balderas, C. et al. Plasma and urine metabolic fingerprinting of type 1 diabetic children. Electrophoresis 34, 2882–2890, doi:10.1002/elps.201300062 (2013).

7 Nelson, R. W. & Reusch, C. E. Animal models of disease: classification and etiology of diabetes in dogs and cats. J Endocrinol 222, T1–9, doi:10.1530/joe-14-0202 (2014).

8 Davison, L. J., Weenink, S. M., Christie, M. R., Herrtage, M. E. & Catchpole, B. Autoantibodies to GAD65 and IA-2 in canine diabetes mellitus. Vet Immunol Immunopathol 126, 83–90, doi:10.1016/j.vetimm.2008.06.016 (2008).

9 Durocher, L. L., Hinchcliff, K. W., DiBartola, S. P. & Johnson, S. E. Acid-base and hormonal abnormalities in dogs with naturally occurring diabetes mellitus. J Am Vet Med Assoc 232, 1310–1320, doi:10.2460/javma.232.9.1310 (2008).

10 O'Kell, A. L. et al. Comparative pathogenesis of autoimmune diabetes in humans, NOD mice, and canines: has a valuable animal model of type 1 diabetes been overlooked? Diabetes (in press) (2017).

11 Lloyd, A. J. et al. Characterisation of the main drivers of intra- and inter-breed variability in the plasma metabolome of dogs. Metabolomics 12, 72, doi:10.1007/s11306-016-0997-6 (2016).

12 Colyer, A., Gilham, M. S., Kamlage, B., Rein, D. & Allaway, D. Identification of intra- and inter-individual metabolite variation in plasma metabolite profiles of cats and dogs. Br J Nutr 106 **Suppl 1**, S146–149, doi:10.1017/s000711451100081x (2011).

13 Minamoto, Y. et al. Alteration of the fecal microbiota and serum metabolite profiles in dogs with idiopathic inflammatory bowel disease. Gut Microbes 6, 33–47, doi:10.1080/19490976.2014.997612 (2015).

14 Li, Q. et al. Veterinary Medicine and Multi-Omics Research for Future Nutrition Targets: Metabolomics and Transcriptomics of the Common Degenerative Mitral Valve Disease in Dogs. Omics 19, 461–470, doi:10.1089/omi.2015.0057 (2015).

15 Patterson, R. E., Ducrocq, A. J., McDougall, D. J., Garrett, T. J. & Yost, R. A. Comparison of blood plasma sample preparation methods for combined LC-MS lipidomics and metabolomics. J Chromatogr B Analyt Technol Biomed Life Sci 1002, 260–266, doi:10.1016/j.jchromb.2015.08.018 (2015).

16 Liu, H., Garrett, T. J., Tayyari, F. & Gu, L. Profiling the metabolome changes caused by cranberry procyanidins in plasma of female rats using (1) H NMR and UHPLC-Q-Orbitrap-HRMS global metabolomics approaches. Mol Nutr Food Res 59, 2107–2118, doi:10.1002/mnfr.201500236 (2015).

17 Pluskal, T., Castillo, S., Villar-Briones, A. & Oresic, M. MZmine 2: modular framework for processing, visualizing, and analyzing mass spectrometry-based molecular profile data. BMC Bioinformatics 11, 395, doi:10.1186/1471-2105-11-395 (2010).

18 Xia, J. & Wishart, D. S. Using MetaboAnalyst 3.0 for Comprehensive Metabolomics Data Analysis. Curr Protoc Bioinformatics 55, 14.10.11–14.10.91, doi:10.1002/cpbi.11 (2016).

19 Lloyd, A. J. et al. Ultra high performance liquid chromatography-high resolution mass spectrometry plasma lipidomics can distinguish between canine breeds despite uncontrolled environmental variability and non-standardized diets. Metabolomics 13, 15, doi:10.1007/s11306-016-1152-0 (2017).

20 Xu, F. et al. Metabolic signature shift in type 2 diabetes mellitus revealed by mass spectrometry-based metabolomics. J Clin Endocrinol Metab 98, E1060–1065, doi:10.1210/jc.2012-4132 (2013).

21 Grapov, D. et al. Diabetes Associated Metabolomic Perturbations in NOD Mice. Metabolomics 11, 425–437, doi:10.1007/s11306-014-0706-2 (2015).

22 Oxenkrug, G., van der Hart, M. & Summergrad, P. Elevated anthranilic acid plasma concentrations in type 1 but not type 2 diabetes mellitus. Integr Mol Med 2, 365–368, doi:10.15761/imm.1000169 (2015).

23 Hipkiss, A. R. et al. Pluripotent protective effects of carnosine, a naturally occurring dipeptide. Ann N Y Acad Sci 854, 37–53 (1998).

24 Janssen, B. et al. Carnosine as a protective factor in diabetic nephropathy: association with a leucine repeat of the carnosinase gene CNDP1. Diabetes 54, 2320–2327 (2005).

25 Steffes, M. W. et al. Diabetic nephropathy in the uninephrectomized dog: microscopic lesions after one year. Kidney Int 21, 721–724 (1982).

26 Jeraj, K., Basgen, J., Hardy, R. M., Osborne, C. A. & Michael, A. F. Immunofluorescence studies of renal basement membranes in dogs with spontaneous diabetes. Am J Vet Res 45, 1162–1165 (1984).

27 Nelson, R. in Canine and Feline Endocrinology (eds EC Feldman, RW Nelson, CE Reusch, & JCR Scott-Moncrief) Ch. 6, 213–257 (Elsevier, 2015).

28 Herring, I. P., Panciera, D. L. & Werre, S. R. Longitudinal prevalence of hypertension, proteinuria, and retinopathy in dogs with spontaneous diabetes mellitus. J Vet Intern Med 28, 488–495, doi:10.1111/jvim.12286 (2014).

29 Wewalka, M., Patti, M. E., Barbato, C., Houten, S. M. & Goldfine, A. B. Fasting serum taurine-conjugated bile acids are elevated in type 2 diabetes and do not change with intensification of insulin. J Clin Endocrinol Metab 99, 1442–1451, doi:10.1210/jc.2013-3367 (2014).

30 Houten, S. M., Watanabe, M. & Auwerx, J. Endocrine functions of bile acids. Embo j 25, 1419–1425, doi:10.1038/sj.emboj.7601049 (2006).

31 Staley, C., Weingarden, A. R., Khoruts, A. & Sadowsky, M. J. Interaction of gut microbiota with bile acid metabolism and its influence on disease states. Appl Microbiol Biotechnol 101, 47–64, doi:10.1007/s00253-016-8006-6 (2017).

32 Tai, N., Wong, F. S. & Wen, L. The role of gut microbiota in the development of type 1, type 2 diabetes mellitus and obesity. Rev Endocr Metab Disord 16, 55–65, doi:10.1007/s11154-015-9309-0 (2015).

